# Thermodynamic principles of enzymatic regulation in biomolecular condensates from reaction-coupled molecular modeling

**DOI:** 10.1101/2025.10.13.682073

**Authors:** Enrico Lavagna, Francesco Delfino, Georgii Koniukov, Matteo Paloni, Luca Ciandrini, Alessandro Barducci

## Abstract

Biomolecular condensates are dynamic cellular assemblies stabilized by weak intermolecular interactions. Cells regulate condensate formation, composition, and function through energy-consuming processes such as post-translational modifications (PTMs), which modify the physicochemical properties of condensate components, thereby dynamically reshaping these interactions. Here, we investigate how enzymatic reactions regulate phase-separated systems using a thermodynamically consistent particle-based model, which allows sampling of out-of-equilibrium steady states. We find that reaction kinetics are intrinsically coupled to the local molecular environment, leading to the formulation of two general principles. First, reactions that weaken favorable interactions are thermodynamically suppressed within condensates. As a consequence, regulation of condensate solubility is most efficient when PTMs tune interactions to values close to the solubility threshold. In contrast, reaction rates are generally enhanced at condensate interfaces, where the thermodynamic inhibition is relieved, but reactant availability remains high. Higher-resolution simulations of FUS and DDX4 proteins indicate that this interfacial contribution remains substantial even for micron-sized condensates, positioning interfaces as key determinants of biochemical activity in phase-separated systems alongside the properties of the condensate bulk. Together, these findings identify general, thermodynamic principles that govern the regulation of biomolecular condensates and link enzymatic activity with phase separation. Moreover, they provide a thermodynamically consistent molecular framework that can be applied to a broad range of regulatory processes in active phase-separated systems.

## Introduction

Cellular condensates are membrane-less organelles composed of proteins and nucleic acids, that take part in many vital processes throughout the nucleus and cytoplasm (1–4). Their assembly and material properties originate from a network of dynamical interactions between one or more condensate components (5–7) that is often modulated by energy-consuming processes, such as enzymatic post-translational modifications (PTMs) acting as fast regulatory switches (8, 9). Such regulatory mechanisms can maintain condensates in non-equilibrium steady states (NESS)(10) which endow them with properties beyond passive compartmentalization (11) and possibly prevent irreversible aggregation(12). Experimental studies have begun to dissect how ATP-dependent reactions shape condensate behavior, with effects ranging from altered morphology and phase boundaries to compositional remodeling and rheological changes (10, 13–16). These findings indicate that active regulation can control condensate properties in ways not captured by equilibrium thermodynamics. Indeed, chemically fueled PTM cycles act on the very intermolecular interactions that sustain phase separation, which in turn modify the local environment in which these reactions take place. This generates a feedback between enzymatic activity and molecular organization which still misses a molecular level understanding. It is unclear, for instance, by which mechanisms phase-separation facilitates or hinders chemical activity, and how these effects can influence the conditions in which PTMs most effectively control condensate solubility. Moreover, the interplay between enzymatic activity and condensate internal structuring remains unknown, despite growing evidence for equilibrium and out-of-equilibrium effects like composition gradients or shell formation (10).

The theoretical community has explored how chemical reactions modulate phase-separated systems using mesoscale continuum descriptions (17–20). These approaches offer insight into how reaction-driven processes influence droplet size, number, and phase boundaries (21). However, their lack of molecular resolution limits their ability to describe key physical mechanisms such as spatial correlations, interaction heterogeneity, and the microscopic organization of components within condensates. Conversely, while molecular simulations are uniquely powerful for investigating the formation, structure, and dynamics of biomolecular condensates (22–25), applying them to systems with chemical reactions remains challenging, particularly when aiming to model reactivity in a thermodynamically consistent way. Early efforts to introduce reactive particle-based models (26, 27) provided valuable insight into the interplay of chemical activity on phase-separated systems but assumed reaction rates that are independent of the local molecular environment, missing the thermodynamic coupling between intermolecular interactions and reaction kinetics. In fact, this assumption neglects the energetic cost of modifying proteins embedded in dense molecular assemblies.

To address these questions, we introduce a particle-based model of a chemically regulated condensate that incorporates reactive dynamics, in which proteins can switch between a phase-separating state and a soluble state, mimicking the action of post-translational modifications. By allowing sustained, energy-consuming conversion between these states, we show how chemical driving shapes condensate composition, spatial organization, and phase behavior under nonequilibrium conditions. In our model, reaction kinetics are coupled to the local molecular environment by enforcing local detailed balance (LDB) (28, 29), that has recently been used to investigate phosphorylation-driven release of TDP-43 from biomolecular condensates (30).

Using this framework, we reveal that phase-separated systems exert a strong thermodynamic feedback on enzymatic activity, leading to an optimal range of PTM modification intensity for maximal solubility control, and to the enhancement of chemical rates at the condensate interface, with consequent substructuring of the phase-separating material. These results identify general principles governing condensate regulation in cells and position interfaces as key determinants of biochemical activity in phase-separated systems, linking reaction kinetics to emergent spatial organization.

## Results

### A Minimal Molecular Model for Biochemical Regulation of Condensates

Here, we introduce a minimal molecular model to investigate the physical basis of condensate regulation by energy-consuming reactions, while circumventing the molecular complexity of real membrane-less organelles. To this end, we adopt an ultra-coarse-grained representation in which each protein is modeled as a single particle. The system includes two molecular species: a scaffold protein that can switch between a condensate-forming (non-phosphorylated, *N*) state and a more soluble (phosphorylated, *P*) state; and a kinase enzyme (*K*) that catalyzes the phosphorylation reaction (*N* + *K* → *P* + *K*).

Interactions between proteins are modeled via Lennard-Jones (LJ) potentials, with the interaction strength and size of the non-phosphorylated scaffold (*ε*_*N*_ = 1, *σ*_*N*_ = 1) defining the energy and length scales. We set the simulation temperature to *k*_*B*_*T* = 0.75, so that the non-phosphorylated scaffold undergoes phase separation with an excess transfer free energy of Δ*G*_trans_ ≃ −4 kJ mol^−1^, consistent with values measured in vitro for model biomolecular condensates (31–33). To model the regulatory effect of phosphorylation, we assign reduced interaction strength to the *P* state *ε*_*P*_ ≲ 0.69, such that they do not phase separate at the chosen temperature. Indeed, since the critical temperature for particles interacting through a truncated and shifted LJ potential with *ε* = 1 is *k*_*B*_*T* = 1.08 (34), it follows that 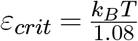 and in our system 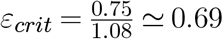. Kinases (*K*) are modeled as particles identical to *N* in terms of interaction strength, so that they co-partition into condensates, in line with *in vitro* (35) *in vivo* (9) observations. All particles are assumed to have the same size (*σ*_*N*_ = *σ*_*P*_ = *σ*_*K*_), and the interaction parameters for all species are summarized in Table S2. The enzymatic reaction can be schematized in two steps. First, we have the formation of the scaffold-enzyme complex, which happens when the two proteins come within a distance of 1.5*σ*. This complex can then react, with a probability dictated by an assigned microscopic rate *λ*_p_. For the sake of simplicity, dephosphorylation is implemented as a unimolecular reaction with rate *λ*_dp_. To reflect the strong directional driving that characterizes both phosphorylation and dephosphorylation in cellular conditions, the corresponding reverse reactions are suppressed by a factor of 10−6. The complete reaction scheme is illustrated in Fig. 1a.

**Figure 1:**
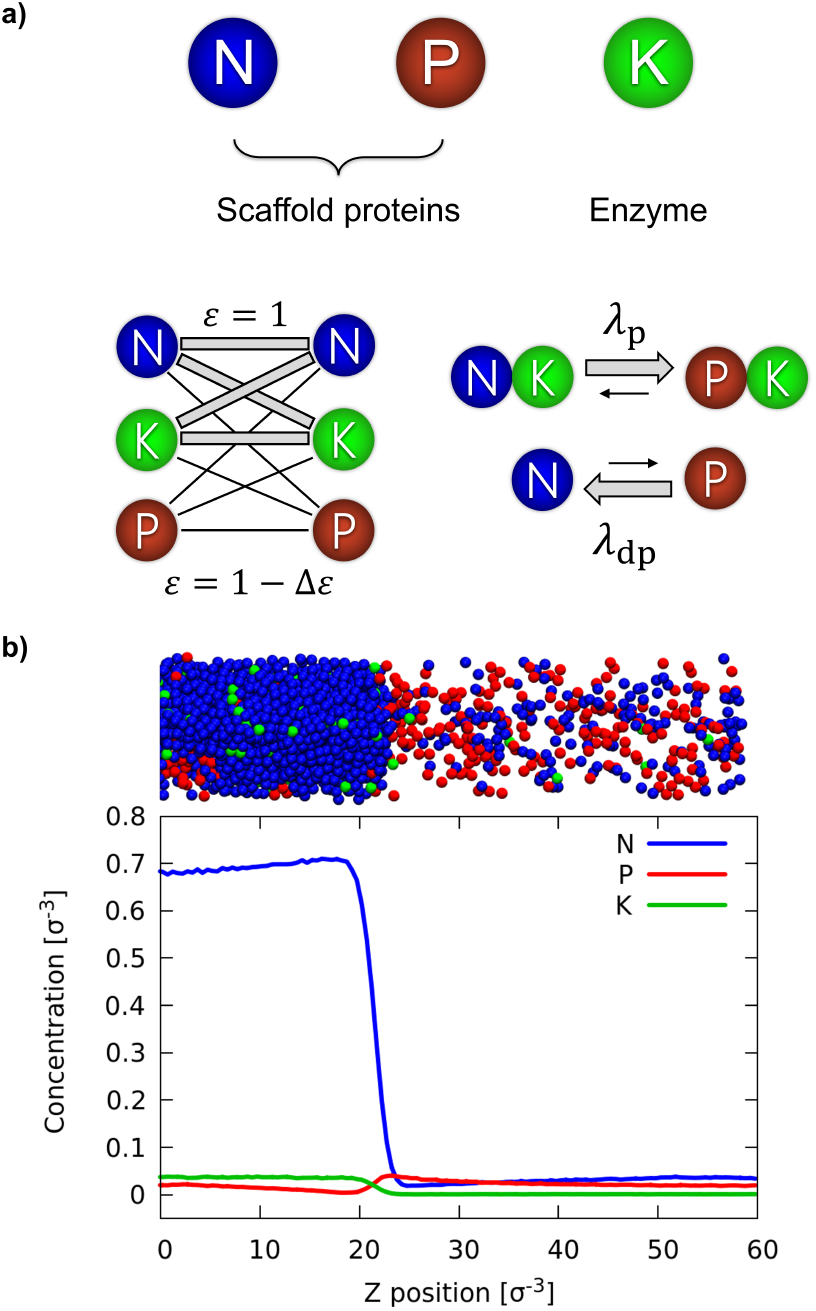
Model and simulation setup. a) Top: the three particles that make up the model. Bottom left: schematics of the interactions. Bottom right: a schematic representation of the implemented reaction cycle. b) Concentration profiles along the z coordinate of a typical slab simulation at steady state, along with a snapshot of the simulated system. Scaffold proteins are depicted in blue, phosphorylated scaffold in red, and protein kinases in green.

Molecular simulations were performed using ReaDDy2 (36), which combines Brownian dynamics for particle motion with Monte Carlo–style steps to model chemical reactions. Each reaction is attempted at a rate *λ* and accepted with a Metropolis-like probability that depends on the change in potential energy associated with the particle transformation. This is crucial in phase separated systems, where the high substrate concentration leads to strong interaction energies. This procedure enforces the principle of local detailed balance (LDB) (28), ensuring thermodynamic consistency by adjusting transition probabilities according to the local environment. Specifically, the probability of a single reaction event is reduced if it would increase the local potential energy. While each reaction obeys LDB, kinase localization introduces an asymmetry between the two reaction pathways, breaking global detailed balance and driving the system into a non-equilibrium steady state.

### Condensate assembly depends on phosphorylation balance

To investigate the interplay between chemical reactions and condensation, we performed phase coexistence simulations using the *slab method*(37–39), in which the system is confined to an elongated box with periodic boundary conditions. This setup, illustrated in Fig. 1b, enables accurate sampling of the dilute and condensed phases as well as the interface, while reducing finite-size effects.

In our reactive system, phase equilibrium depends on the competition between phosphorylation and dephosphorylation processes. We therefore explore the system behavior by varying the phosphorylation microscopic rate *λ*_p_, keeping the dephosphorylation rate fixed at *λ*_dp_ = 0.005 *τ*^−1^. We note that the results presented here are valid for a wide range of *λ*_p_ and *λ*_dp_ values, as better detailed in the SI (Fig. S3b). For this set of simulations, we fixed *ε*_*P*_ = 0.3, so that phosphorylated proteins are effectively soluble. After a transient, the system reaches a steady state for all values of *λ*_p_, with higher values leading to a larger fraction of phosphorylated proteins *N*_*P*_ (Fig. S4a).

We characterize the spatial organization in the resulting stationary states by constructing a phase diagram as a function of *λ*_p_ using dilute and condensed phase concentration of all proteins in the system (Fig. 2a). As the reaction cycle becomes more biased toward phosphorylation, the coexistence region progressively narrows: the dilute-phase concentration rises with *λ*_p_, i.e., pushing the reaction balance towards phosphorylation. At the same time, the condensed-phase concentration remains largely unchanged. At *λ*_p_ ≥ 1.0 *τ*^−1^ the condensate dissolves completely, and the system becomes homogeneous. This is due to the compositional effect of increasing *λ*_p_: as more scaffold proteins are phosphorylated, they become more soluble and are depleted from the condensed phase, resulting in its gradual shrinkage. Since the total number of proteins is fixed, the increase in the scaffold protein number *N*_*P*_ leads to an enrichment of the dilute phase. The resulting phase behavior closely matches that of a passive, non-reactive system with fixed composition, where the fraction *N*_*P*_ */N*_*T*_ (where *N*_*T*_ is the total protein number) directly controls phase separation (Fig. S4b).

**Figure 2:**
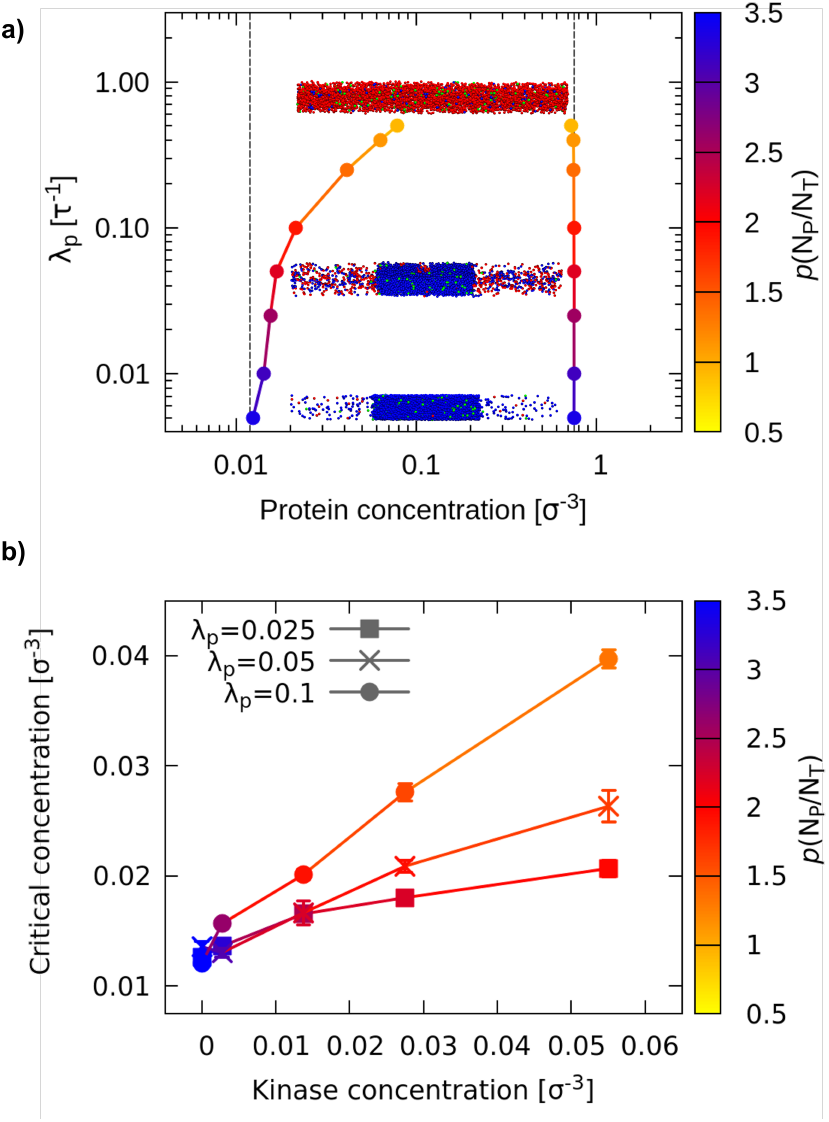
Phase separation depends on reaction imbalance and enzyme concentration. a) Phase diagram as a function of *λ*_p_ at fixed kinase concentration. For each value of *λ*_p_, the two points represent the total protein concentration in the dilute and condensed phases, respectively. Each point is shaded to show the ratio of P particles (*N*_*P*_) over the total (*N*_*T*_) at steady state, following the scale on the left. The dashed bars represent the dilute and condensed phase concentrations at equilibrium. The snapshots illustrate different product concentrations *P* at different *λ*_p_ points and how they correlate with phase separation. b) Saturation concentration as a function of kinase concentration, for three fixed values of *λ*_p_ .

We then explore how the phase behavior depends on the concentration of kinases *ρ*_*K*_. Figure 2b shows the saturation concentration (*i*.*e*. the dilute-phase boundary) as a function of *ρ*_*K*_, revealing a linear dependence that mirrors experimental observations for the DYRK3 kinase (9). Consistent with the trend observed when varying *λ*_p_, this behavior reflects the fact that increasing the number of kinases raises the steady-state fraction of phosphorylated proteins *N*_*P*_ */N*_*T*_, which, in turn, governs the degree of condensation. Overall, these results show that enzymatic activity controls condensate assembly primarily by tuning the steady-state fraction of phase-separating proteins, effectively acting as a non-equilibrium control parameter for phase coexistence.

### Modification strength maximizes regulatory efficiency at the solubility threshold

PTMs can alter the energy of intermolecular interactions to various degrees, especially when considering the possibility of multi-site modifications (40–42). A recent study suggests that multisite phosphorylation provides a tunable mechanism for controlling protein-protein interaction strength: only after a critical number of sites have been phosphorylated, the homotypic interaction is weakened enough for the proteins to leave the condensed phase (30). Here, we investigate how such variability in PTM-induced interaction modifications affects the steady-state properties of phase separation. We model this feature of a PTM-driven system by introducing the parameter

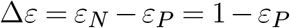

which quantifies the change in intermolecular interaction energy upon enzymatic modification. We refer to this property as the *modification strength*. An increase in Δ*ε* corresponds to a stronger phosphorylation effect, which disfavors the interaction between phosphorylated proteins and the rest of the system.

Interestingly, the simulations reveal a non-monotonic relationship between the phase separation propensity of the system and the modification strength, as illustrated by the snapshots in Fig. 3a. We quantify the dependence of condensation on the modification strength Δ*ε* by tracking the total partition coefficient *C*_p_ of the system (Fig. 3b). *C*_p_ is the ratio of the protein concentrations in the condensed (*ρ*_*dens*_) and dilute phase (*ρ*_*dil*_): higher values of *C*_p_ mean that proteins favor partitioning inside the condensate. For small modification strengths (Δ*ε* ≤ 0.3), increasing Δ*ε* leads to a decrease in *C*_p_, i.e., phase separation becomes less favorable. This trend is coherent with our expectation of enhanced phase separation regulation, as higher Δ*ε* values favor the solubilization of *P* in the dilute phase. When increasing Δ*ε* beyond 0.3, however, *C*_p_ increases: yet again, phase separation tends to be favored. Hence, *C*_p_ has a minimum at Δ*ε* ≈ 0.3, which means that the system is at maximum solubility. The snapshots visually highlight this effect, showing a significantly denser dilute phase for the simulation at Δ*ε* = 0.3. Decoupling the effect on *C*_p_ of the concentrations of the two phases (Fig. 3c), we observe that the non-monotonic behavior is mostly due to the variation in dilute concentration, with the condensed phase concentration remaining mostly constant. The complete phase diagrams at different values of Δ*ε* (Fig. S5b) further show that the system has the narrowest phase separation region at Δ*ε* ≈ 0.3 . Notably, this value corresponds to the interaction energy below which *P* proteins are unable to phase separate on their own. Furthermore, this non-monotonic behavior is more pronounced for higher values of the phosphorylation rate, *λ*_p_ (Fig. S5a).

**Figure 3:**
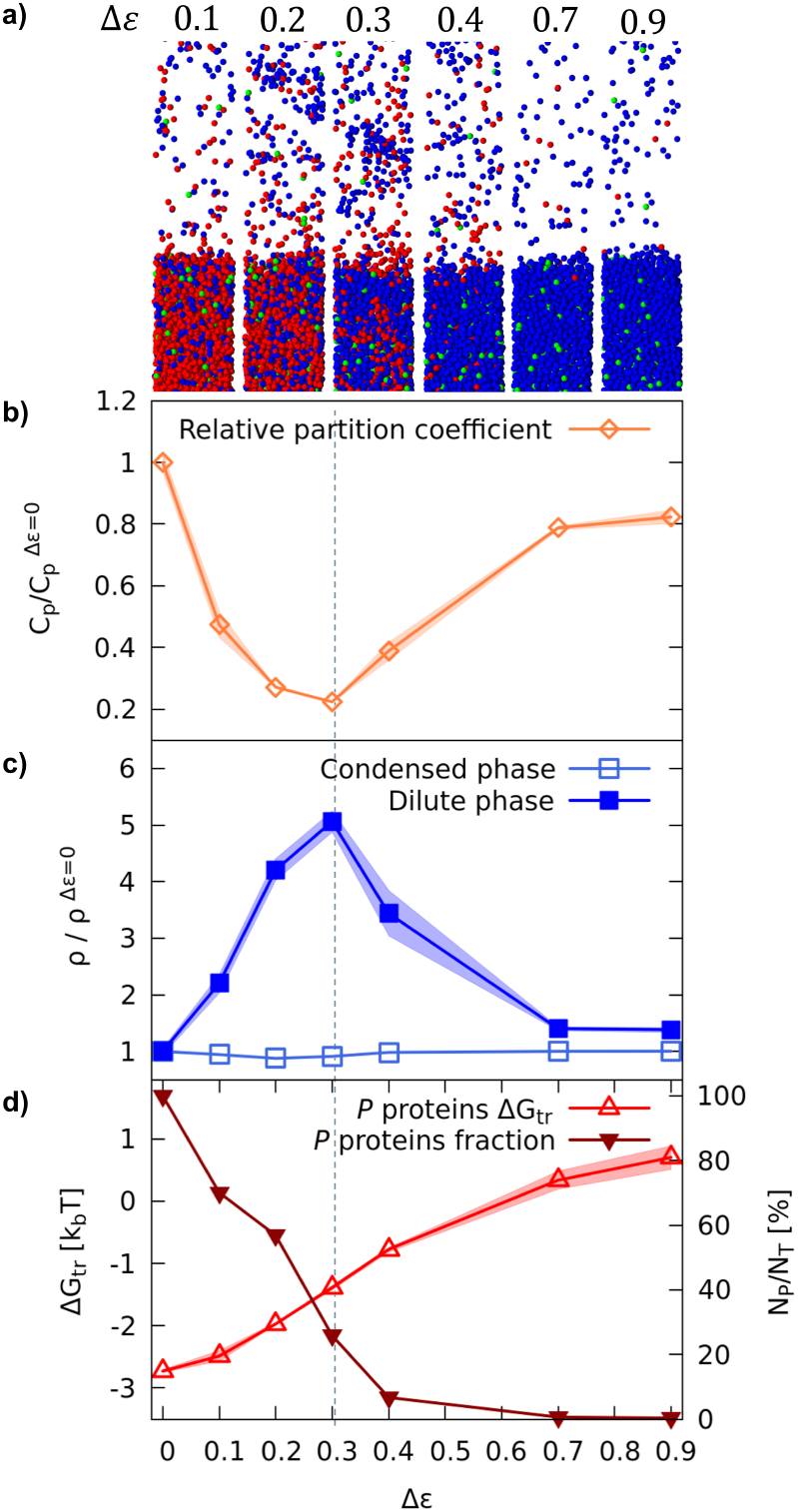
Phase behavior depends on the intensity of interaction modifications induced by phosphorylation. a) Snapshots from the simulations showing the phase coexistence. a) Partition coefficient of all proteins as a function of Δ*ε*, normalized with respect to its value at Δ*ε* = 0. Lower *C*_p_ means that proteins prefer the dilute phase, which increases the critical concentration. The grey line represents the critical Δ*ε* over which *P* proteins are no longer able to phase separate on their own. c) Protein concentration of the dilute phase (full squares) and dilute phase (empty squares), normalized with respect to their value at Δ*ε* = 0. d) Free energy of transfer of *P* proteins compared to their fraction as a function of Δ*ε*. Higher transfer free energies mean that proteins prefer to partition in the dilute phase (see Methods). The point at Δ*ε* = 0 is taken from the simulations with no reactions and only scaffold proteins. All data presented in this figure was obtained from simulations at *λ*_p_ = 0.05.

To rationalize this behavior, we recall that phase separation is primarily determined by the steady-state *P* protein fraction, as established in the previous section. In Fig. 3d, we report the dependence of the *P* protein fraction (*N*_*P*_ */N*_*T*_) and the transfer free energy (Δ*G*_trans_) of *P* proteins between the two phases on Δ*ε*. As expected, Δ*G*_trans_ increases with Δ*ε*, indicating that *P* proteins increasingly disperse in the dilute phase. Concurrently, however, the fraction of *P* proteins decreases drastically. This reduction in *P* fraction compensates for their increased solubility, explaining why the condensed phase is stabilized at high Δ*ε*: although individual *P* proteins are more soluble, their lower abundance diminishes their overall disruptive effect on phase separation. Furthermore, the height of the *P* fraction curve increases with *λ*_p_*/λ*_dp_, which explains the higher dilute concentrations at higher ratios, as Δ*G*_trans_ itself shows no such dependency (Fig. S5c).

The drop in *P* protein fraction arises because stronger modification imposes a higher energetic penalty for the phosphorylation reaction. This ultimately leads to a lower phosphorylation rate, particularly within the densely packed condensed phase. For Δ*ε* ≤ 0.3, the *P* fraction remains high, but their attraction to unmodified *N* proteins is still sufficient to retain them in the condensed phase, as seen in the two leftmost snapshots of Fig. 3. Regulation is thus most effective near the solubility threshold of *P* proteins. At this point, the modification is sufficient to induce their dispersion from the condensate, yet their concentration remains high enough to significantly hinder phase separation. More generally, these findings define an optimal energetic regime for active control, in which post-translational modifications tune condensate stability by placing the system at the edge of phase separation.

### The condensate interface is a nexus of chemical activity

So far, we have characterized the system behavior in terms of global observables, such as phase diagrams and average protein concentrations. We now explore local properties to better understand the interplay between chemical regulation and spatial organization, particularly at the interface.

We first examine the spatial distribution of phosphorylation and dephosphorylation reaction fluxes, calculated as the total amount of reaction per unit of time and surface across the *xy* plane (Fig. 4a). Remarkably, the phosphorylation flux exhibits a pronounced peak at the interface between the condensed and dilute phases. Notably, this spatial localization of chemical activity is not expected in models that assume homogeneous reaction rates, and instead emerges from the local modulation of reaction kinetics by the local environment. Inside the condensate, the high concentrations of scaffold proteins (*N*) and kinases (*K*) favor frequent encounters between enzymes and substrates. Still, the dense environment also increases the energetic penalty for converting *N* to *P*, which reduces the effective reaction rate (Figure 4b top panel). This suppression is modulated by the modification strength Δ*ε*: stronger modifications incur a larger energetic penalty, further reducing phosphorylation rates in the condensed phase (Fig. S6a), consistent with the non-monotonic behavior of phase stability described in the previous section. At the interface, *N* and *K* concentrations remain relatively high, but scaffold proteins experience fewer stabilizing interactions, lowering the energetic cost of phosphorylation. This leads to enhanced reaction activity specifically at the boundary between phases. In contrast to phosphorylation, the dephosphorylation flux closely follows the spatial distribution of *P* proteins. Since converting *P* to *N* generally leads to more favorable interactions, the reaction does not incur an energetic penalty. As a result, its effective rate is largely insensitive to the local environment, and the flux is primarily shaped by substrate availability.

**Figure 4:**
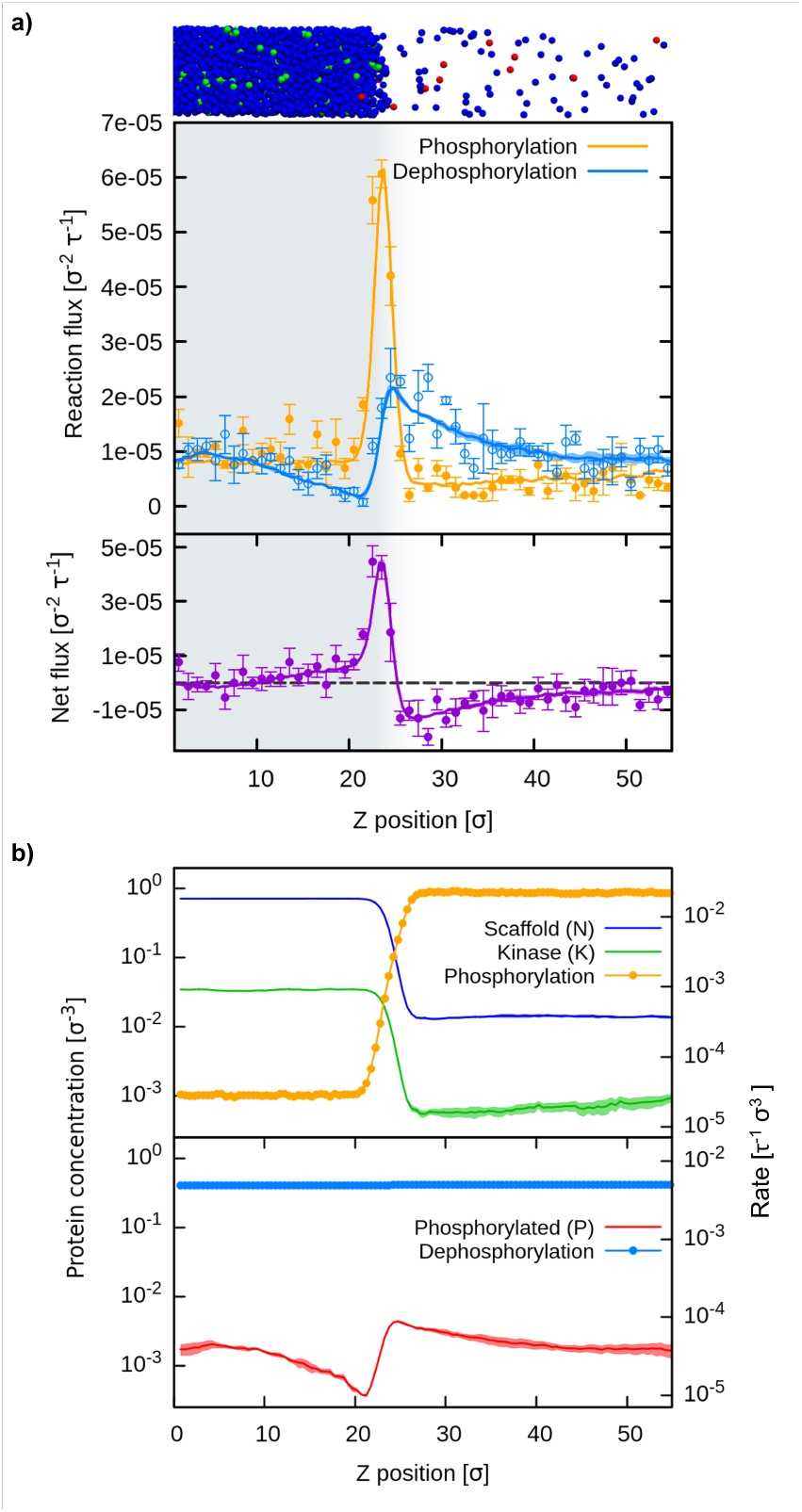
Phosphorylation accumulation at the condensate’s interface. In this figure, we represent different quantities as a function of the distance from the center of the condensed phase. Data from the simulations at *λ*_p_ = 0.05 *τ*^−1^, *ϵ*_*P*_ = 0.3. a) Rate flux of the two reactions. Fluxes were quantified by direct reaction counting (dots), and reaction activity recomputation (line, see Methods). The bottom plot shows the net particle flux at the interface (positive means more *P* proteins produced). The shaded area indicates the extension of the condensed phase. b) Comparison between protein concentration and microscopic reaction rates of phosphorylation (top panel) and dephosphorylation (bottom panel). Protein concentrations are plotted on the left y-axis, while the microscopic rate is plotted on the right y-axis.

The phosphorylation and dephosphorylation fluxes do not compensate near the phase boundary, resulting in a net production of phosphorylated proteins (*P*) within the condensed phase and of non-phosphorylated proteins (*N*) in the dilute phase (Fig. 4a, bottom panel). This asymmetric reaction distribution gives rise to a localized and directional flux across the interface with *P* proteins continuously exiting the condensate while *N* proteins are replenished from the dilute phase. Such sustained directional fluxes are a hallmark of non-equilibrium systems driven by active processes. To assess the impact of the interface also under equilibrium conditions, we simulate a version of the model with no explicit enzymes in which both reactions are reversible and unimolecular. As expected, the system shows no net chemical flux. Nonetheless, both reactions display pronounced peaks near the interface, showing that spatial variations in density and interactions can concentrate reaction activity even in the absence of energy input (Fig. S6b).

To corroborate the potential role of the interface as a localized catalytic environment, the question arises whether in real-world biomolecular condensates the interfacial region is large enough that its relative weight can significantly impact the overall chemical activity. Indeed, assuming that the width of the interface only depends on the interaction of the condensate components and not on its size, the ratio of the interface and total volumes will decrease with the condensate radius. Hence, we perform slab simulations of phase-separating disordered domains from the FUS and DDX4 proteins (FUS-LC and NDDX4, respectively) to measure the spatial extension of the interface, based on the intermolecular energy profile (see Methods section). Assuming a fixed interface width *l* = 14 nm, as measured in these higher resolution simulations (Fig. 5a), we can then calculate the weight of the interface chemical activity (Fig. 5b). This percentage can be quite high (*>* 20%) even for spherical condensates larger than 1 μm. Since the size of typical cellular condensates rarely exceeds the 2 μm diameter, we note that the weight of the interface flux is significant even for very large condensates. Considering that many cellular biomolecular condensates are smaller than 1 μm and may exhibit non-spherical shapes, we deem this a conservative estimate of the impact of interfacial catalytic enhancement.

**Figure 5:**
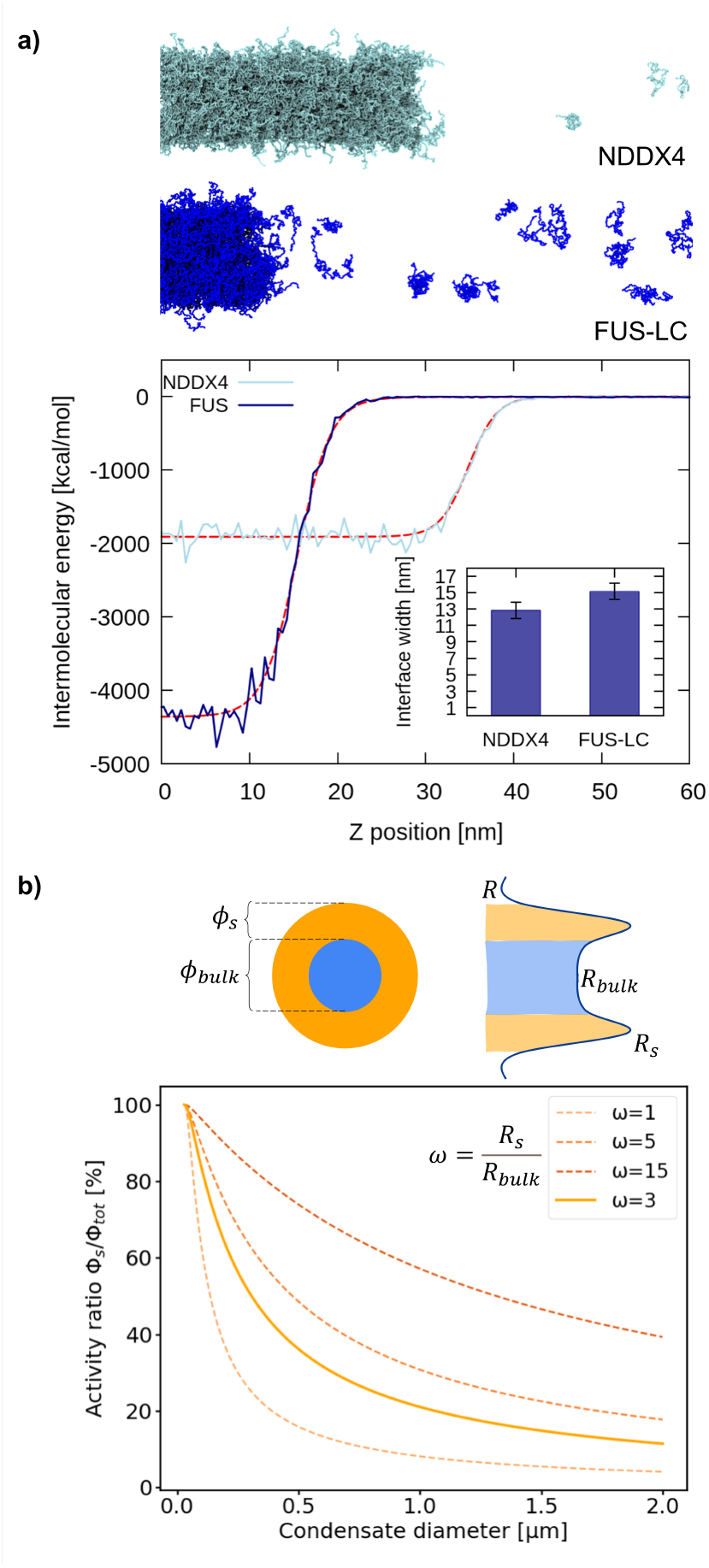
The interface effect is relevant for real-world condensates. a) Intermolecular energy as a function of the distance from the slab center, obtained from the CALVADOS simulations of NDDX4 and FUS-LC protein domains. The dashed red lines represent the fitted data, through which we extracted the interfacial width shown in the inset (see Methods section). On top, snapshots of the phase coexistence simulations of the two systems. b) Estimate of the relative weight of the interfacial activity in a spherical condensate, as a function of the diameter. In the sketch, we illustrate the involved quantities: the total amount of reactions per time Φ_S_ and Φ_bulk_ are obtained by multiplying the average rates *R*_S_ and *R*_bulk_ by the respective volumes. The ratio Φ_S_*/*Φ_tot_ is plotted at fixed interfacial width *l* = 14 nm and at different values of *ω*. These values have the same order of magnitude as the ratio *R*_*S*_*/R*_bulk_ measured in our simulations (see Methods).

## Discussion

In this study, we identify general physical principles governing the phase behavior and reactive properties of biomolecular condensates regulated by enzymatic modifications. With our model, we are able to show how enzymatic activity modulates the nonequilibrium phase behavior of the system: as the balance shifts toward phosphorylation, either through an increased kinase rate or a higher kinase concentration, biomolecular condensates are increasingly solubilized, and ultimately dissolve (19). In our simulations, enzymatic regulation plays a role analogous to temperature in upper critical solution temperature (UCST) systems, acting as a control parameter for condensed phase stability. The presence of an intrinsic nonequilibrium drive leads to a broadened spectrum of possible phase behaviors, which we characterize by constructing the phase diagram as a function of the phosphorylation/dephosphorylation imbalance. Here, we provide such a description at the molecular level, allowing us to capture different mechanisms that depend on the local dynamics and interactions of proteins and enzymes. This molecular resolution reveals a reciprocal coupling between enzymatic activity and phase separation, whereby reactions shape condensate organization and condensates reshape reaction kinetics. In particular, two key results arise from this coupling.

First, we identify a principle for optimizing regulation by tuning the affinity between scaffold proteins and those modified by PTMs. Condensate solubilization depends non monotonically on the modification strength of interparticle interactions. At fixed enzyme concentration and catalytic rates, regulation is most effective when the modification lowers the intermolecular interaction just below the phase-separating threshold. Beyond that threshold, conversion inside the condensed phase becomes increasingly costly, which suppresses production of the modified state and weakens the overall solubilizing effect. This principle suggests a simple design rule for synthetic or engineered condensate systems in which the phase separating material is produced or suppressed via chemical reactions (13): efficient control can be achieved with reaction cycles that only modestly reduce the affinity between condensate components. More broadly, this principle suggests that effective condensate regulation is achieved when modification places the system close to the solubility threshold. This is in line with the idea that cellular proteins have evolved to be near the edge of solubility, where small changes in molecular interactions can produce large functional responses (43, 44).

This effect arises because our simulations show a local modulation of reaction activity, which depends not only on the local concentrations of reactants, but also on their state in terms of interaction energy. As a direct consequence, reactions that weaken favorable interactions between condensate components are suppressed in the bulk of the condensed phase, where the energetic penalty is more severe. This energetic penalty, on the other hand, is significantly reduced in the interfacial region, where the reactant concentration is still high enough to favor substrate-enzyme encounters.

This leads to the second principle identified by our simulations, namely the dramatic enhancement of enzymatic activity at the condensate interface. Higher-resolution simulations of representative phase-separating systems (FUS-LC and NDDX4) corroborate this finding by showing that the interfacial region represents a significant fraction of the total condensate volume. The interface then becomes a new, major determinant in how phase separation regulates biochemical activity, along with crowding and localization effects (45). Previous studies (19, 46) proposed that a positive phosphorylation flux inside the condensate, which scales volumetrically as *R*^3^, would compensate Ostwald ripening, which scales as *R*, for a critical radius. Even if enzymatic activity were entirely concentrated at the interface, the reaction flux would scale as *R*^2^, which is still sufficient to stabilize condensate size. In this direction, there is already evidence showing enhanced reaction rates in smaller enzymatic condensates (47), suggesting that activity may scale with surface rather than volume, consistent with the hypothesis of enhanced interfacial activity. We note that the interfacial effect on chemical reactions arises whenever, in an active phase-separating system, the condensed phase is enriched in reactants and reduced in products. While this situation is typical of enzymatic regulation (19, 30, 35), it can also occur in a wide range of biochemical processes, both enzymatic and non-enzymatic (Fig. S6), for which condensates are known to act as catalytic platforms (48). Moreover, the increased interface activity leads to product accumulation at the interface, which may provide a mechanistic basis for experimentally observed features of chemically driven steady-state condensates, such as core-shell substructuring (10, 49). While the functional relevance of condensate interfaces has only recently been investigated (50, 51), our results reveal a new mechanism whereby crowding-modulated reaction rates drive interface structuring.

In summary, our simulations reveal key features of chemically regulated cellular condensates, with minimal assumptions on the biological nature of the involved proteins. We identify an optimal range of interaction modifications for active regulation and highlight the central role of the condensate interface. We show that these phenomena emerge naturally when the proper thermodynamic constraints are used to couple reaction kinetics to the local environment.

While the high level of coarse-graining in our model prevents the investigation of finer effects that PTMs can have on protein-protein interactions, the mechanisms identified here only rely on generic thermodynamic constraints and enzymatic driving. Therefore, the principles we define are broadly relevant for chemically regulated biomolecular condensates. Building on this generality, the model provides a foundation for exploring a broader range of systems characterized by different reaction schemes or condensate architectures, paving the way towards a better understanding of how enzymatic regulation shapes the organization and function of biomolecular condensates.

## Materials and Methods

### 1-bead-per-molecule simulations

To simulate the steady state of our phase-separating ultra-CG model, we relied on the ReaDDy2 software package (36). The time evolution of particle diffusion was simulated following the overdamped Langevin equation, with the diffusion coefficient *D* for all molecules set to unity. These simulations were run with a time-step 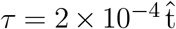, with 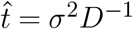 being the time unit in our reduced system, as better detailed in section Extended Methods of the SI. Particles interacted through a truncated and shifted Lennard-Jones pair potential, with the truncation set at the standard value of 2.5*σ*. The reactions were handled by the LDB algorithm included in ReaDDy (28). At each time step and for each reaction, all reactive particles would attempt a reaction step with probability *P* = 1 − *e*^−*λτ*^, with *λ* the microscopic rate for that reaction. After that, the reaction was accepted or rejected with a Monte Carlo check of the system’s potential energy. We choose microscopic reaction rates of the driven reactions in the range of 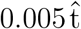 to 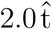, which allowed for proper sampling of the phase separation steady-state. A more thorough discussion on the connection between microscopic rates in our model and real-world enzymatic systems can be found in section of the SI.

All phase coexistence simulations were performed in triclinic boxes with periodic boundary conditions of size *x* = *y* = 11 *σ* and *z* = 120 *σ*, with the condensed phase spanning the *xy* plane. The boxes were initialized with the particles arranged in a cubic lattice with minimal distance *σ*, placed in the center of the box to obtain a slab configuration. Each simulation contained 4400 particles. We let the simulations run until both the number of particles and the concentration of dilute and condensed phases reached a steady state for at least 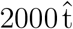. Since equilibration times varied depending on initial configuration, composition, and set of parameters, not all simulations had the same run-time, but all ranged between 3000 and 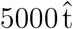. All data, including system trajectory, were saved with a frequency of 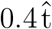 (2000 *τ*). We replicated each simulation three times, and steady-state quantities are measured as the average over these three replicas.

For the phase diagram in *λ*_p_, we explored *λ*_p_ values by varying the phosphorylation rate *λ*_p_ while keeping *λ*_dp_ = 0.005 *τ*^−1^, for each value of Δ*ε*. We fixed the kinase concentration at*ρ*_*K*_ = 0.014 *σ*^−3^, low enough to prevent phase separation by kinases themselves.

### Reactions

The chemical reactions included in the model are the kinase-mediated phosphorylation and a spontaneous dephosphorylation (we implicitly consider the phosphatase to be diffuse in the system). These reactions are schematized below, with particles not explicitly included in the system written within round brackets

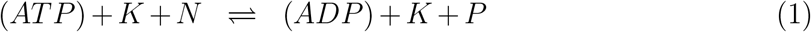

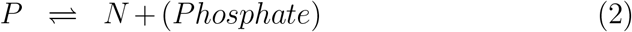

We label the forward reaction rates of equation 1, 2, i.e. reading the reactions from left to right, respectively 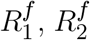, while the backward rates 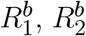 correspond to reading reactions 1, 2 from right to left.

Overall conversion rates *N* → *P* (*R*_*N*→*P*_) and *P* → *N* (*R*_*P* →*N*_) are thus defined as

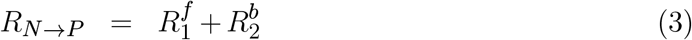

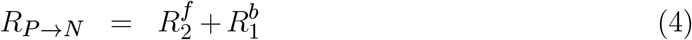

In the dilute limit, reaction rates can be written as follows

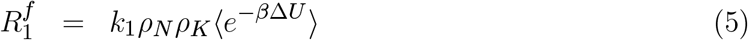

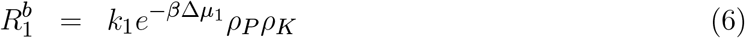

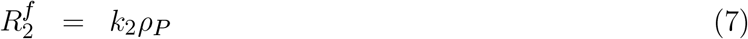

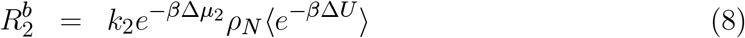

where *k*_1_ and *k*_2_ are microscopic transition probabilities, *β* is the inverse of *k*_*B*_*T, ρ*_*i*_ is the concentration of species *i* and ⟨*e*^−*β*Δ*U*^⟩ represents the ensemble average of the Boltzmann factor used in the LDB algorithm. Backwards rates are suppressed by a factor 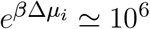 with respect to their corresponding forward rates, which takes into account the chemical driving of implicit particles involved in reaction *i*. In our reaction scheme, Δ*μ*_1_ = *μ*_*AT P*_ − *μ*_*ADP*_ and 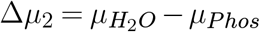.

### 1-bead-per-residue simulations

Slab simulations of FUS-LC and NDDX4 protein condensates were performed using the OpenMM package with the CALVADOS2 force field (52). The simulations were produced using a Langevin integrator with time step 0.01 ps at a temperature of 300 K and a relaxation time of 100 ps. All simulations were carried out in two parts: an equilibration in the NPT ensemble to stabilize the condensed phase and a production run in the slab configuration in the NVT ensemble. Total simulated time in the production run was 10 μs.

### Calculation of dilute and condensed phase concentrations

In a slab simulation, properties like the concentration of particles can be plotted in a dome-like profile, as visible in Fig. 1a. To extract the concentrations of condensed and dilute phases from these configurations, we first computed the total density profiles, averaging over 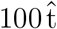 time windows and symmetrizing with respect to the center of the condensate, identified with the position of the center of mass of the system. Then, for each profile, we performed a fit based on an adapted hyperbolic tangent function, as proposed by Tesei et al. (53).

### Calculation of the free energy of transfer

To calculate the free energy of transfer, Δ*G*_trans_, we exploited the results from the fit of the concentration with the hyperbolic tangent function to set the *z* position of the interfaces between the two phases. For each frame, we counted the number of protein molecules within the condensed phase, *N*_in_, and the dilute phase, *N*_out_. The partitioning coefficient was determined as the ratio of the respective number densities, *K*_*p*_ = (⟨*N*_in_⟩*/V*_in_)*/*(⟨*N*_out_⟩*/V*_out_), where *V*_in_ and *V*_out_ are the volumes of the two phases. The free energy was then obtained from the relation Δ*G*_trans_ = −*k*_*B*_*T* ln *K*_*p*_.

### Recomputation of expected local rates

To calculate the local reaction rates without relying only on reaction counts, which are relatively low, we performed an indirect recomputation based on the trajectory. In each step of the saved trajectory, we looped over all reactive particles and measured the Δ*U* associated with the available reaction. From the Δ*U*, we could obtain the acceptance probability *a*(Δ*U*) and the reaction rate of that particle, and average their values over all particles in the same z bin and over time. Since the rates obtained through the recomputation are in good agreement with the direct reaction count and have relatively lower signal noise, we utilized those to illustrate other analyses throughout this study, including acceptance probability and Δ*U* .

### Calculation of the interface flux percentage

While the difference in intermolecular energy inside and outside the condensed phase varies significantly between condensates composed of the two proteins, the width of the interface is similar and measures around 14 nm (Fig. 5a). We estimate the relative weight of the reaction flux for a spherical condensate as a function of its radius, using this equation

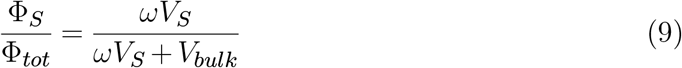

where Φ_*S*_ and Φ_*tot*_ are the total reaction fluxes from the interface and the whole condensate, 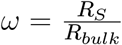 is the ratio between the average reaction flux in the interface region and bulk of the condensate, *V*_*S*_ the volume of the interface and *V*_*bulk*_ the volume of the bulk (see the sketch in Fig. 5b). From this equation, it is possible to express 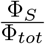 as a function of the condensate radius and width of the interface shell.

## Acknowledgments

We acknowledge the support of the Swiss National Science Foundation under Grant CRSII5 193740 and of the French Agence Nationale de la Recherche under grant ANR-21-CE30-0001 and Projet ANR-21-CE12-0023.

## Author contributions

E.L. contributed to model development, performed simulations, analyzed data and interpreted results. F.D. contributed to model development and preliminary simulations. G.K. performed 1-bead-per-residue simulations and contributed data analysis. M.P. contributed to data analysis and interpretation. L.C. contributed to model development and interpretation. A.B. conceived and supervised the investigation, interpreting results and directing all contributions. E.L. and A.B wrote the manuscript, with contributions from all authors for editing and review.

