## Supplementary materials for "Thermodynamic principles of enzymatic regulation in biomolecular condensates from reaction-coupled molecular modeling"

Table S1: Units of measure in the simulations.

| Unit | Natural | DYRK3 |
|---|---|---|
| Energy | $\epsilon$ | |
| Length | $\sigma$ | $5\ nm$ |
| Time | $\sigma^2 D^{-1}$ | $10^{-5} s$ |

### Supplementary materials

#### Extended Methods

##### 1-bead per molecule simulations

Our reactive model is defined by the presence of two different particles, the enzyme $K$ and the scaffold protein, which can change its state from non-phosphorylated ($N$) to phosphorylated ($P$) and vice versa, following the reaction scheme described in Fig 1a, including two main reaction paths and their inverse. Hence, each simulated system is fully characterized by the following parameters: interaction energy of the non-phosphorylated ($\epsilon_P$), microscopic reaction rates ($\lambda_{\text{p}}$, $\lambda_{\text{dp}}$) and particle densities ($\rho_S$, $\rho_K$).

**Units** We define our units of measure in terms of the properties of the scaffold protein $N$, i.e. $\epsilon_{NN}$, we set $\sigma_{NN}$ and $D_N$ to unity and all other quantities are measured relative to them. Thus, any observable can be expressed in common unit if we assign to each of these reference quantities a reasonable real-world value. For example the kinase DYRK3 has $\sim 588$ residues and $MM \simeq 67\,\text{kDa}$. Its radius and diffusion coefficient could be approximately $R \sim 2.5\,\text{nm}$ and $D \sim 2.5\,\mu\text{m}^2\,\text{s}^{-1}$. Table S1 reports a possible mapping of the reduced units in our model to common units as obtained from comparison with DYRK3 characteristics features.

**Algorithm** We performed molecular dynamics simulations relying on a hybrid simulation protocol, which combined Brownian dynamics in the continuous configurational space and the possibility for the molecules to have transitions between discrete biochemical states. Brownian motion dynamics is obtained from the numerical solution of the overdamped Langevin equation:

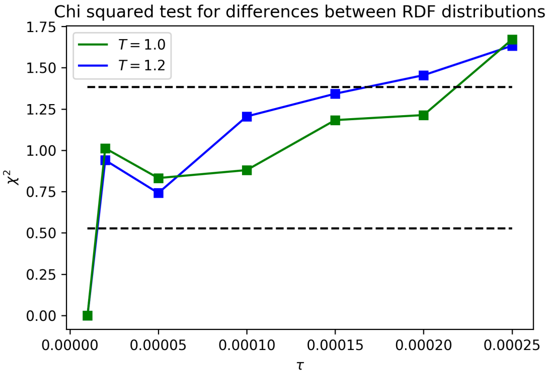


Figure S1: Reduced chi squared test applied to radial distribution functions computed at different values of the timestep for a system in the liquid phase with $\rho = 0.7\sigma^{-3}$. We consider the timestep $\tau = 10^{-5}\sigma^2/D$ as a reference. The two dashed lines represent respectively a 99.5% probability and a 5% probability of obtaining greater values for the chi squared.

$$\dot{\mathbf{r}}(t) = -D\frac{\nabla U(\mathbf{r}(t))}{k_B T} + \sqrt{2D}\boldsymbol{\xi}(t)$$

where $D$ is the diffusion coefficient and $\boldsymbol{\eta}(t)$ is white noise with $\langle \eta_i(t)\eta_j(t')\rangle = 2D\delta_{ij}\delta(t-t')$. The integrator solves the equation numerically using a Euler-Maruyama discretization.

Local detailed balance in the time evolution is ensured by a Metropolis-like Monte Carlo step in the reaction handling algorithm. Local detailed balance ensures that the reaction probability correctly takes into account the local (around the reaction site) environment configuration, i.e. transitions to energetically disfavoured configurations are suppressed by the customary Boltzmann factor. The algorithm was described in ref (28).

**Timestep** We want to set the simulation timestep as large as possible without incurring in numerical errors during the simulation. To probe the best value for the timestep we run simulations for systems with only particles $N$ without reactions for different values of the temperature and the density. We compare radial distribution functions (RDF) obtained for various values of the timestep with the RDF computed with a reference timestep $\tau = 10^{-5}\sigma^2/D$. We observe that there are no significant differences for timestep up to $2 * 10^{-4}\sigma^2/D$, see Fig.S1. Thus the timestep of all our simulations is set to $2 * 10^{-4}\sigma^2/D$.

Table S2: Interactions between particle species.

| | N | P | K |
|---|---|---|---|
| **N** | 1 | $1-\Delta\varepsilon$ | 1 |
| **P** | | $1-\Delta\varepsilon$ | $1-\Delta\varepsilon$ |
| **K** | | | 1 |

**Interactions** All interactions are modeled by the following functional form, a truncated and shifted Lennard-Jones potential. The shift is defined by imposing the potential equal to zero at the cutoff.

$$V(r) = \begin{cases} V_{LJ}(r) - V_{LJ}(r_{cut}) & r < r_{cut} \\ 0 & r > r_{cut} \end{cases} \tag{10}$$

with

$$V_{LJ}(r) = 4\epsilon\left(\left(\frac{\sigma}{r}\right)^{12} - \left(\frac{\sigma}{r}\right)^{6}\right) \tag{11}$$

Each protein-protein pair interaction is characterized by the $\varepsilon$ parameter, which in our systems has two possible values: unity if the interaction involves the non-phosphorylated scaffold, and $\varepsilon_P = 1 - \Delta\varepsilon$ if it involves the phosphorylated scaffold. These interactions are summarized in table S2. For the protein to lose ability to phase separate, $\varepsilon_P$ should be $\lesssim 0.69$ with our choice of temperature (34) (Illustrated in Fig. S2).

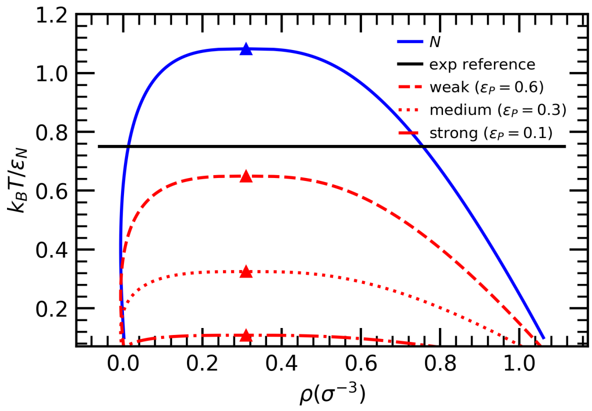


Figure S2: Sketch of the temperature dependence of the LJ binodal curve in our shifted and truncated LJ model.

**Diffusion coefficient** The diffusion coefficient of proteins in cells is approximately $D \sim 0.1$ to $10\,\mu m^2\,s^{-1}$ (54). In *E. coli* the diffusion coefficient $D$ ($\mu m^2\,s^{-1}$) as a function of the molecular mass $MM$ (kDa, where $1\,Da = m(C^{12})/12$) can be estimated by the following expression:

$$D = \alpha(MM)^{-2} + D_0 \tag{12}$$

with $\alpha = 4.3 \times 10^3\,\mu m^2\,s^{-1}\,kDa^2$ and $D_0 = 0.65\,\mu m^2\,s^{-1}$ (55).

**1-bead per residue simulations**

**Protein sequences** Using Calvados parameters, we prepared our configurations using the following protein sequences:

- **NDDX4**:
  MGDEDWEAEINPHMSSYVPIFE
  KDRYSGENGDNFNRTPASSSEM
  DDGPSRRDHFMKSGFASGRNFG
  NRDAGECNKRDNTSTMGGFGVG
  KSFGNRGFSNSRFEDGDSSGFW
  RESSNDCEDNPTRNRGFSKRGG
  YRDGNNSEASGPYRRGGRGSFR
  GCRGGFGLGSPNNDLDPDECMQ
  RTGGLFGSRRPVLSGTGNGDTS
  QSRSGSGSERGGYKGLNEEVIT
  GSGKNSWKSEAEGGES

- **FUS-LC**:
  MASNDYTQQATQSYGAYPTQPG
  QGYSQQSSQPYGQQSYSGYSQS
  TDTSGYGQSSYSSYGQSQNTGY
  GTQSTPQGYGSTGGYGSSQSSQ
  SSYGQQSSYPGYGQQPAPSSTS
  GSYGSSSQSSSYGQPQSGSYSQ

QPSYGGQQQSYGQQQSYNPPQG

YGQQNQYNS

**Initial configuration** For each simulation, we used the gmx insert-molecules tool to insert 400 copies of the protein molecules, in random positions and orientations, in $25 \times 25 \times 500\,\mathrm{nm}^3$ boxes. Then the NPT ensemble simulation was carried in order to naturally form the tight dense phase with a shorter z-length ( 30 nm).

#### Reaction rates

In our simulation algorithm macroscopic reaction rates $R_i^j$ (5, 6, 7, 8) result from microscopic rates $r_i^j$, which are expressed in terms of a reaction probability $P(\lambda_i^j)$ per particle per timestep $\tau$

$$\lambda_i^j = \frac{P(r_i^j)}{\tau} \tag{13}$$

$P(r_i^j)$ is the product of two factors, a proposal probability $p$ and an acceptance probability $a$. The proposal probability $p$ depends only on the constant microscopic rate $\lambda$

$$p = 1 - e^{-\lambda\tau} \simeq \lambda\tau \tag{14}$$

$\lambda$, which contains all the microscopic details, is treated here as an input parameter that can be tuned to obtain the desired macroscopic rate.

The acceptance probability $a$ depends on the energy difference between the proposed state and the starting state, according to the Metropolis algorithm standard acceptance function

$$a = min\left\{1, \frac{V_{reac,N}^{eff}}{V_{reac,P}^{eff}} e^{-\beta\Delta U}\right\} \tag{15}$$

where $V_{reac,N/P}^{eff}$ is the effective reaction volume, defined by

$$V_{reac,N/P}^{eff} = \int_0^{R_{reac}} e^{-\beta U_{A/B}(r)} 4\pi r^2 dr \tag{16}$$

with $U_{N/P}(r)$ the interaction potential between $N/P$ and the enzyme $K$. The Boltzmann factor, with $\Delta U = E_P - E_N > 0$ the difference in the interaction energy of the protein with the environment, suppresses transitions to higher energy configurations. Expressions 5, 6, 7, 8 satisfy local detailed balance conditions

$$\frac{R_1^f}{R_1^b} = \frac{\rho_N}{\rho_P} e^{-\beta \Delta U} e^{\beta \Delta \mu_1} = e^{\beta(\mu_N - \mu_P + \Delta \mu_1)} \tag{17}$$

$$\frac{R_2^f}{R_2^b} = \frac{\rho_P}{\rho_N} e^{\beta \Delta U} e^{\beta \Delta \mu_2} = e^{\beta(\mu_P - \mu_N + \Delta \mu_2)} \tag{18}$$

where we used the definition of chemical potential $\mu_i = E_i + k_B T \ln \rho_i$. Inserting 14 and 15 into 13 we obtain the expression of the microscopic rates given that reagents have been already brought together. In the case of a conversion reaction like 2 reaction events are evaluated for each particle in the system, thus macroscopic rates 7, 8 are simply proportional to the particle concentration and

$$\lambda_{\mathrm{dp}}^f = k_2 \tag{19}$$

$$\lambda_{\mathrm{dp}}^b = k_2 e^{-\beta \Delta \mu_2} \tag{20}$$

In the case of an enzymatic reaction like 1 a reaction event occurs by definition if the distance between reagents is below a reaction radius $R_{reac} = 1.5\sigma$. In sufficiently dilute conditions, i.e. as long as the law of mass action holds, macroscopic rates 5, 6 are proportional to the product of the concentrations of reagents, with the proportionality constant $k_1$ having units of per time per concentration. In this case the relation between $k_1$ and $\lambda_{\mathrm{p}}$ involves also taking into account the probability that reagents come in contact

$$\lambda_{\mathrm{p}}^f \simeq \frac{k_1}{V_{reac,N}^{eff}} \tag{21}$$

$$\lambda_{\mathrm{p}}^b \simeq \frac{k_1 e^{-\beta \Delta \mu_1}}{V_{reac,P}^{eff}} \tag{22}$$

**Inverse reaction rates** Each of the two main reaction pathways, phopshorylation and dephosphorylation, is modeled with an associate inverse reaction for thermodynamic consistency. Inverse rates are determined by the Gibbs free energy difference associated to the reactions in physiological conditions. Measured values of Gibbs free energy ($\Delta G^0$)

Table S3: Gibbs energy released in the hydrolysis of different types of phosphate bonds. Physiological values are obtained by considering the following concentrations: $[ATP] \simeq 3\ mM$, $[ADP] \simeq 0.6\ mM$ and $[P] \simeq 10\ mM$.

| **reaction** | $\Delta G^0$ | $\Delta G^{phys}$ |
|---|---|---|
| $ATP + H_2O \rightarrow ADP + P$ | 13 $k_BT$ | 20 $k_BT$ |
| serine phosphate $+H_2O \rightarrow$ serine $+P$ | 10 $k_BT$ | 15 $k_BT$ |
| threonine phosphate $+H_2O \rightarrow$ threonine $+P$ | 8 $k_BT$ | 13 $k_BT$ |
| tyrosine phosphate $+H_2O \rightarrow$ tyrosine $+P$ | 12 $k_BT$ | 17 $k_BT$ |

for hydrolysis reactions in standard conditions are reported in table S3.

Free energies obtained in standard conditions need to be corrected to take into account physiological concentrations of ATP, ADP, and phosphate. For a generic estimate, we consider physiological concentrations to be: $[ATP] \simeq 3\ mM$, $[ADP] \simeq 0.6\ mM$, and $[P] \simeq 10\ mM$. The Gibbs free energy in physiological conditions is then

$$\Delta G^{phys} = \Delta G^0 + k_BT \ln Q \tag{23}$$

The corrective term $Q$ is equal to the ratio of products concentrations over reactants concentrations expressed with physiological concentrations (normalized by standard concentrations, i.e. $[1M]$). Calculated values of $\Delta G^{phys}$ are reported in the last column of table S3. The Gibbs free energy for the phosphorylation reaction (1) can be obtained from the difference between the free energy of ATP hydrolysis and that of phosphate ester hydrolysis. Given the energy differences between different types of phosphate esters, we consider their average weighted with their respective frequency in human phosphorylated protein sites (86% Serine, 12% Threonine, 2% Tyrosine). Resulting Gibbs free energy differences are of $5k_BT$ for the phosphorylation (1) and of $15k_BT$ for the dephosphorylation (2), corresponding to an inverse rate suppression factor respectively of $\gtrsim 100$ and $\gtrsim 10^6$. In the end, to ensure proper irreversibility in our simulations, we chose a common suppression factor of $10^6$ for both reaction pathways.

**Estimation of phosphorylation rate** Expressions 5, 6 are derived in the dilute limit approximation, where the law of mass action holds. The local detailed balance simulation protocol however ensures that expressions 5, 6 hold in scenarios far from the dilute limit, such as the droplet interior, where the rates cannot be easily described by means of analytical expressions. A macroscopic description of enzyme kinetics is usually given in terms of the Michaelis-Menten kinetic model, which allow us to provide a quantitative interpretation of our microscopic rates. In this framework the rate of product formation for a reaction $E+S \rightarrow E+P$ is given by

$$\frac{d[P]}{dt} = k_{cat}\,[E]\,\frac{[S]}{K_M+[S]} \tag{24}$$

where the constant $K_M$ is numerically equal to the substrate concentration at which the reaction rate is half of its maximum value $k_{cat}\,[E]$. The Michaelis-Menten catalytic rate $k_{cat}$ is the maximum number of substrate molecules converted to product per enzyme molecule per second. The model is valid under the assumption that enzyme concentration is small compared to substrate concentration or $K_M$: $[E] \ll K_M+[S]$.

The $k_{cat}$ values of the enzymes in our simulations can be assumed to be of the same order of magnitude of the microscopic rate $\lambda_{\mathrm{p}}$, which ranges between 0.005 and $2\,\sigma^{-2}D$. Converting this quantity using the assumed values of $\sigma$ and $D$ reported in Tab. S1, we obtain $k_{cat} \approx 10^4$ to $10^6 s^{-1}$.

#### Nonequilibrium effects

Chemical regulation of cellular condensates, as explained in the introduction, relies on the nonequilibrium nature of the cytoplasmic or nuclear environment. Enzymatic reactions like phosphorylation and other PTMs require a steady source of energy in the form of a chemical fuel, for instance, ATP. In our simulations, even if the fuel is modeled implicitly, the system is out of chemical equilibrium thanks to the enzymatic reaction cycle. With such a scheme, which also depends on the proximity of the reactants, the global detailed balance of transition between chemical states is intrinsically broken. For instance, in our case, the transition $K+N \rightarrow K+P$ is heavily favored compared to the transition $N \rightarrow P$, and the opposite is true for the inverse reactions.

The non-equilibrium nature of our systems manifests in some of the condensate features, like the diffusive flux at the interface shown in Fig. 4a. Moreover, the concentration

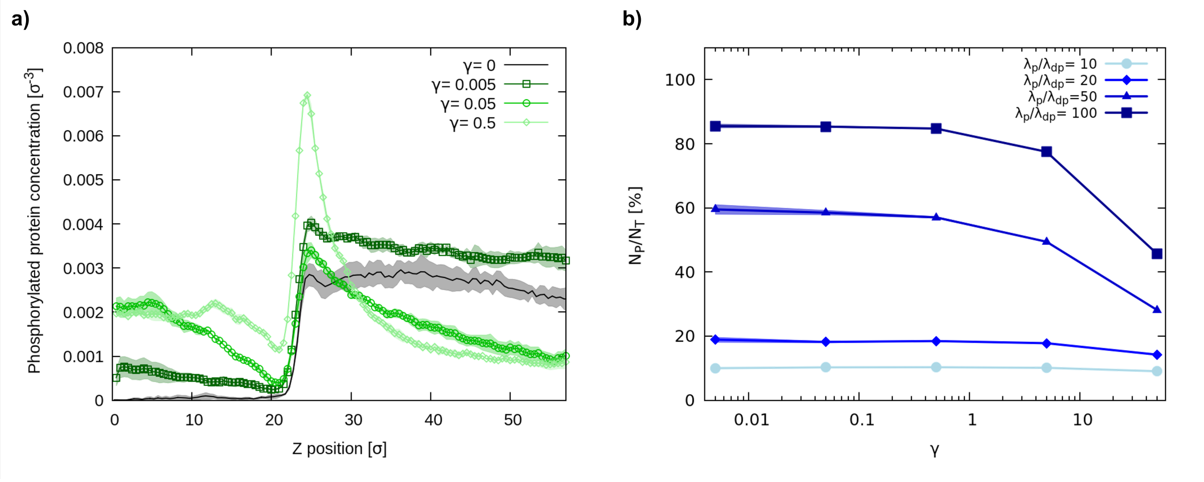


Figure S3: a) Concentration profiles of the phosphorylated protein. The black line is obtained from a set of simulations with no reactions and a fixed $P$ concentration of $0.0017\sigma^{-3}$. b) Total percentage of phosphorylated proteins in small box simulations as a function of $\gamma$.

profile of the $P$ protein is affected, showing a double cusp shape around the condensate interface. To better probe this effect, we performed simulations varying the strength of the non-equilibrium drive, which we quantify with how fast the reactions are catalyzed compared to the diffusive time scale of the simulation. For this quantity, we used the ratio $\gamma = \frac{\lambda_\text{p}}{D} =$ where $\lambda_\text{p}$ is the microscopic rate of phosphorylation (the higher between those of the two reaction paths) and $D$ is the diffusion coefficient in the Langevin equation, which in our simulation is always set to 1.

In Fig. S3a, we show that the shape of the profiles is indeed dependent on $\gamma$, with the relaxation length increasing with lower values of $\gamma$. The profile at $\gamma = 0$ is obtained from a simulation with the same model and total concentration but no reactions, with the $\rho_P$ fixed to a value close to that reached in the other three plotted simulations. At equilibrium, the $P$ protein is almost completely depleted from inside the condensate. Moreover, the asymptotic values of the $P$ concentration (i.e., in the bulk of the condensed phase and in the dilute phase far from the interface) are different in the presence of reactions compared to equilibrium simulations. However, in the simulations at $\gamma = 0.05$ and 0.5 they tend to the same value. In the simulation at $\gamma = 0.005$, the concentration varies more slowly and does not reach an asymptotic value within our simulation boundaries. Its interface features are thus more similar to those at equilibrium. In summary, increasing the rates compared to diffusion leads to stronger but more confined diffusive fluxes.

In the main text, we explain how all of the novel results derive from the steady-state relative concentration of phosphorylated protein, $N_P/N_T$. We investigated how this quantity changes with the intensity of the non-equilibrium by exploring $\gamma$ values comprised between 0.01 and 100 in ad hoc, small box simulations. The resulting values of the relative concentration of phosphorylated protein are shown in Fig. S3. For large enough values of $\gamma$, the $N_P/N_T$ decreases. This results from the different nature of the two reactions: phosphorylation, being enzymatic, is diffusion-limited, while dephosphorylation is unimolecular. Thus, large values of $\gamma$ favor dephosphorylation. For a wide range of $\gamma$ values, however, the concentration is constant, starting to decrease for $\gamma > 1$.

While characteristic catalysis rate for many real world phosphorylation reactions is around $k_{cat} \simeq 10^{-2} - 100\ s^{-1}$ (54, 56), the $k_{cat}$ values resulting from the microscopic rates in our simulations go up to $\simeq 10^6$, as explained in the previous section. Therefore, our rates tend to be an upward estimate of real ones, a choice that enhances the sampling of the steady-state. Nevertheless, in all the slab simulations used for this work, we selected values of microscopic rates $\lambda$ such that $\gamma$ is always $\ll 1$: in this regime, $N_P/N_T$ is constant. Therefore, even if the simulated systems do show signature nonequilibrium features, the presented results are true regardless of the non-equilibrium intensity.

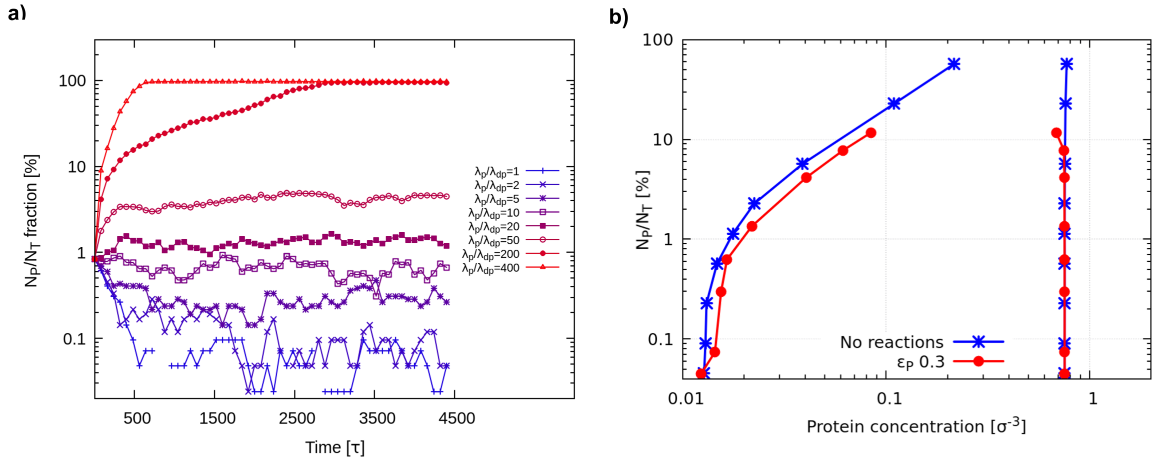


Figure S4: a) Time evolution of the number of phosphorylated proteins in the simulation at $\lambda_\mathrm{p} = 0.05\,\tau^{-1}$, $\varepsilon_P = 0.3$, the ones used for the results in Fig. 2. b) Phase diagrams as a function of $N_P/N_T$. In blue, the equilibrium simulations, where the $N_P/N_T$ concentration is fixed from the beginning; in red, the active system simulated at $\lambda_\mathrm{p} = 0.05\,\tau^{-1}$, $\Delta\varepsilon = 0.7$.

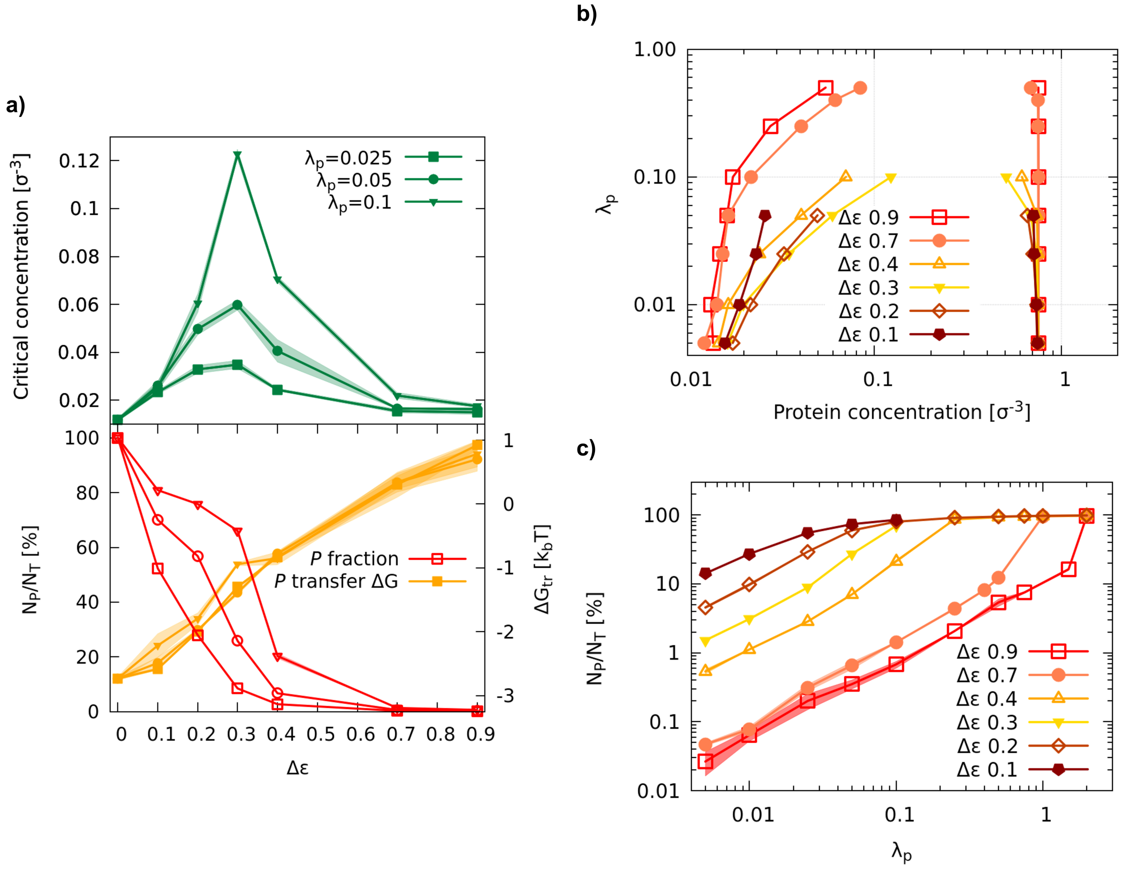


Figure S5: a) Critical concentrations (top) and corresponding $P$ fraction and $\Delta G_{trans}$ (bottom) as a function of $\Delta\varepsilon$, for $\lambda_{\mathrm{p}} = 0.025\,\tau^{-1}$, $0.05\,\tau^{-1}$, and $0.1\,\tau^{-1}$. b) Phase diagram as a function of $\lambda_{\mathrm{p}}$ and for different values of $\Delta\varepsilon$.Critical concentrations as a function of $\Delta\varepsilon$, for $\lambda_{\mathrm{p}} = 0.025\,\tau^{-1}$, $0.05\,\tau^{-1}$, and $0.1\,\tau^{-1}$. c) Phosphorylated protein relative concentration $N_P/N_T$ as a function of $\lambda_{\mathrm{p}}$, for all $\Delta\varepsilon$ values.

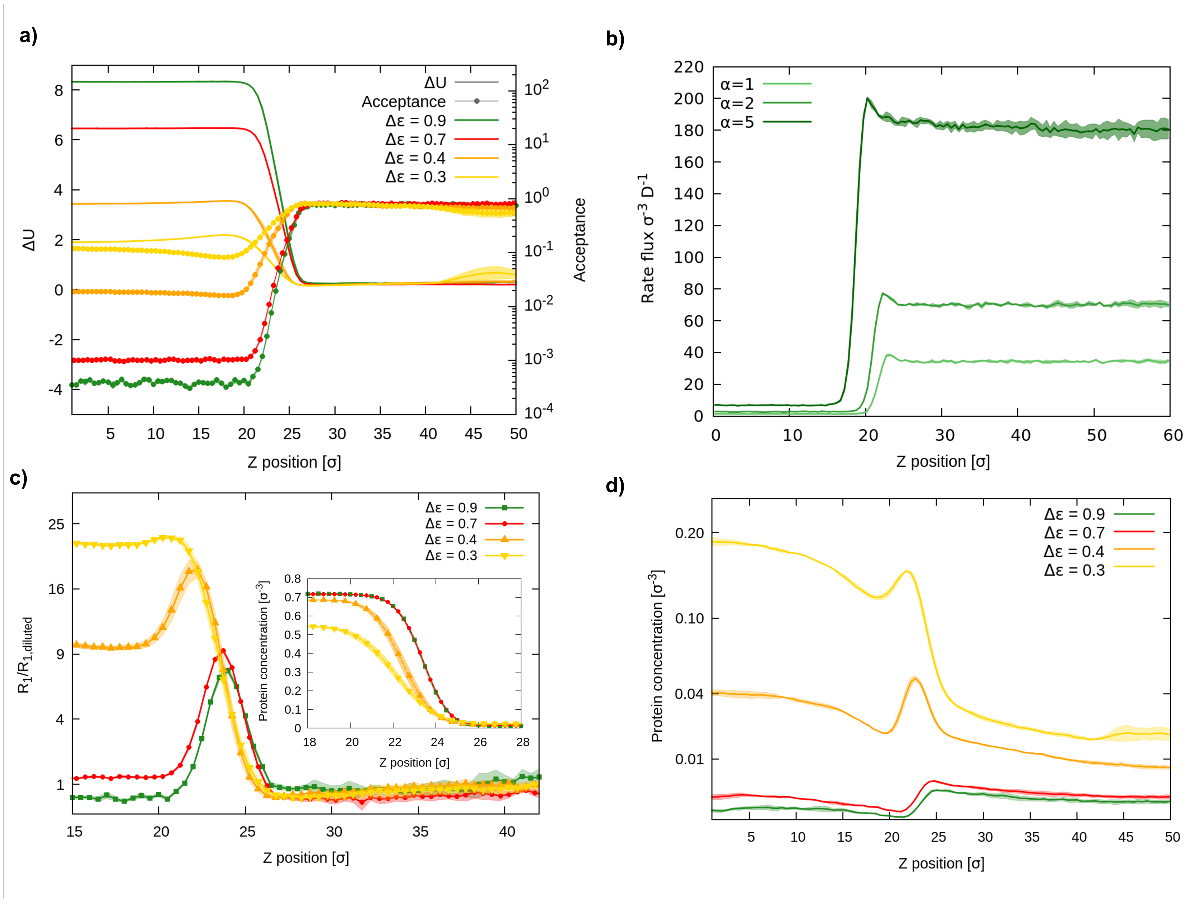


Figure S6: a) $\Delta U$ and acceptance for the phosphorylation reaction as a function of the distance from the center of the condensed phase, for $\Delta\varepsilon = 0.1$, 0.2, 0.6, and 0.7. Outside the condensate, $\Delta U \simeq 0$ and the acceptance is almost 1. b) Rate flux of the simulations with no enzymes. Since the simulation is at chemical equilibrium, this rate flux is the same for both phosphorylation and dephosphorylation. At the interface, we observe a peak in the rates, corresponding to an accumulation of phosphorylated protein $P$. The three lines represent simulations with different values of the parameter $\alpha$, which is the ratio between the microscopic rates of phosphorylation and dephosphorylation. c) Ratio between the phosphorylation reaction flux and its value in the dilute phase, distant from the interface, for different values of $\Delta\varepsilon$. In the inset, the concentration of $N$ proteins, which mark the position and width of the interface for the respective simulations. d) $P$ protein concentration profile at different values of $\Delta\varepsilon$, showing the accumulation of product at the interface.
